# Acoustically-Targeted Measurement of Transgene Expression in the Brain

**DOI:** 10.1101/2023.05.23.541868

**Authors:** Joon Pyung Seo, James S. Trippett, Zhimin Huang, Ryan Z. Wang, Sangsin Lee, Jerzy O. Szablowski

## Abstract

Gene expression is a critical component of brain physiology and activity, but monitoring this expression in the living brain represents a significant challenge. Here, we introduce a new paradigm called Recovery of Markers through InSonation (REMIS) for noninvasive measurement of gene expression in the brain with cell-type, spatial, and temporal specificity. Our approach relies on engineered protein markers that are designed to be expressed in neurons and exit into the interstitium. By applying ultrasound to targeted brain regions, these markers are released into the bloodstream, where they can be readily detected using biochemical techniques. REMIS can noninvasively confirm gene delivery and measure endogenous signaling in specific brain sites through a simple insonation and a subsequent blood test. Using REMIS, we successfully measured chemogenetic induction of neuronal activity in ultrasound-targeted brain regions. REMIS recovery of markers is reliable and demonstrated improved recovery of markers from the brain into the blood in every tested animal. Overall, our work establishes a noninvasive, spatially-specific means of monitoring gene delivery outcomes and endogenous signaling in mammalian brains, opening up possibilities for brain research and noninvasive monitoring of gene therapies in the brain.

## INTRODUCTION

Monitoring gene expression in the living brain is crucial for studying its network activity^1^, diagnosing neurological diseases from onset to progression^2^, and translating gene therapies into the clinic. However, mapping brain gene expression poses significant challenges, especially within the deep tissue. Currently, to directly localize gene activity in deep brain regions, tissue-destructive techniques such as biopsy and post-mortem histology are used^3^. Although common, these approaches prohibit repeated measurements, thus precluding longitudinal assessment of the same subject. Further, they hamper translational and neuroscientific studies, where damage to the brain region being investigated introduces a scientific confound.

Several noninvasive imaging tools have emerged to detect endogenous gene expression within the brain. These include magnetic resonance imaging (MRI) with genetically-encoded contrast agents^4-6^, positron emission tomography (PET) using transgene-binding probes^7^, optical imaging^8-11^, and biomolecular ultrasound^12^. However, these tools must overcome significant barriers, including sensitivity constraints, the need for brain accessible radioactive probes, poor access to deep tissues, and restricted penetration through the skull, respectively. These obstacles have greatly hindered our ability to measure with spatial precision the endogenous expression of critical neuronal activity markers, including immediate early gene *c-Fos*. For example, while MRI-based strategies hold such promise, they currently lack the signal levels necessary for effective *c-Fos* detection. More sensitive techniques like intravital imaging with luciferases can measure *c-Fos* activity in small animals^13^, but light generated by luciferases is subject to tissue scattering and skull attenuation limits use cases in deep brain regions or large animals and prevents accurate localization of the source^14^.

Recently, focused ultrasound-mediated blood-brain barrier opening^15^ (**FUS-BBBO**) was shown to release proteins from the brain^16, 17^ into the circulation, in a process called FUS liquid biopsy. However, this process can only release naturally-existing tissue markers, preventing measurement of many physiological processes in the cells for which secreted markers do not exist, and relying on the naturally existing concentrations of these markers which may be difficult to detect.

We have recently pioneered the concept of synthetic markers of gene expression that originate from the brain and cross through the intact blood-brain barrier (BBB) using the process of reverse transcytosis^18^. These markers, called Released Markers of Activity, or **RMA**^19^ originate from the genetically-labeled cells in the brain and can inform on transduction or endogenous promoter activity through a simple blood test. RMAs are highly sensitive, with the ability to monitor as few as approximately 800 cells, and enable multiplexed monitoring of gene expression. While sensitive, convenient, and noninvasive, the readout of RMAs lacks spatial precision and instead reports on the entirety of RMA-labeled population.

Here, we present Recovery of Markers through InSonation (**REMIS**) as a sensitive, noninvasive alternative to existing methods for site-specific gene expression measurement in the brain. Under this approach, designer markers are expressed within the brain in response to gene activity, and then released from targeted brain regions into the blood via focused ultrasound application. Our strategy incorporates FUS-BBBO to noninvasively target specific brain regions with millimeter precision, enabling the transport of synthetic markers from the brain into blood. From there, the markers can be detected using conventional biochemical assays such as enzyme-linked immunosorbent assay^20^, detection of luciferases^14^, and mass spectrometry^21, 22^, facilitating routine measurements of gene expression from blood draws.

To establish REMIS (**Fig. 1**), we first expressed markers in neurons under constitutive promoters and confirmed noninvasive transduction at targeted brain regions. Next, we reliably quantified the signals of the markers in the blood following FUS-BBBO. Lastly, we implemented the markers to measure endogenous neuronal signaling activity. Specifically, we expressed them under the control of a genetic circuit that responds to c-Fos when activated by heightened neuronal activity^23^. We were able to measure the corresponding neuronal activity of the targeted brain regions through blood tests, a process which typically cannot be measured with blood sampling. Overall, our work demonstrates the feasibility of combining genetically-encoded reporters and focused ultrasound to noninvasively and specifically measure endogenous gene expression in the brain.

**Figure 1.**
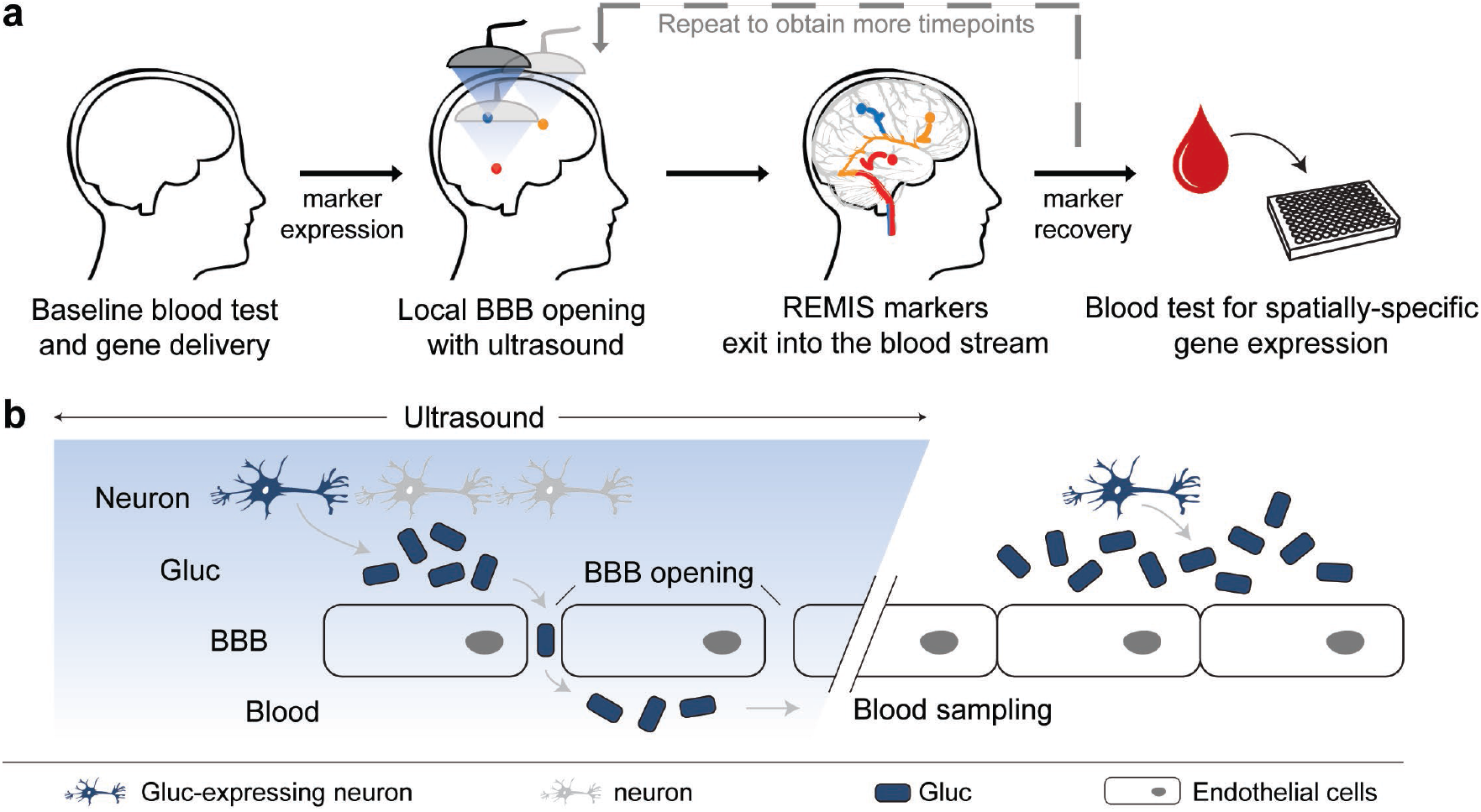
The concept of noninvasive Recovery of Markers through Insonation (REMIS). (**a)** REMIS uses focused ultrasound to open the blood-brain barrier (FUS-BBBO) and release genetically-encoded synthetic markers from the insonated brain regions into the blood. The markers are then collected from simple blood draw for further analysis. Marker levels in the serum can inform on the transgene expression or endogenous physiologic activity in the targeted brain region. **(b)** Any protein that is secreted from the cell can be used as REMIS marker. In the presence of the ultrasound-opened BBB, these secreted markers produced in transduced cells diffuse into the blood. Since ultrasound can be focused with millimeter precision, the markers released into the blood come from a spatially-defined brain region. Overall, REMIS process enables noninvasive, spatially-specific monitoring of genetically-targeted cells in the brain through a simple blood test.

## RESULTS

### Secreted protein reporter as a spatially-specific marker for gene expression in the brain

Our REMIS platform relies on engineered protein markers that get expressed in cells and then released into the brain parenchyma. After FUS is used to open the BBB at specific target sites^24^, the markers are released into the blood, where they can be detected using biochemical methods (**Fig. 1**). In principle, any serum-detectable protein that can be secreted from cells can be used with REMIS. As our readout, we chose *Gaussia* luciferase (GLuc) for the reporter protein. GLuc is a highly sensitive reporter that emits bioluminescence through an enzymatic reaction. It is also naturally secreted from the cell as it harbors a cell secretion signal peptide^25^.

To monitor the neuronal activity in specific brain regions, we constructed an adeno-associated viral (AAV) vector expressing GLuc under the control of the Synapsin 1 (hSyn1) promoter, which drives specific transgene expression in neurons. As an initial proof-of-concept, we delivered the hSyn1-GLuc vector to the entire mouse brain using a BBB-permeable, brain-specific AAV serotype called PHP.eB^26,27^ (GLuc-AAV) and evaluated subsequent GLuc levels in the blood after FUS-BBBO insonation (**Fig. 2a**). Here, we intravenously (i.v.) injected mice with GLuc-AAV at 8.3×10^9^ viral particles per g of body weight following the dose from our previous studies^24, 28^. Afterwards, we measured GLuc bioluminescence levels while the BBB was still intact to obtain baseline GLuc signal levels. Next, we insonated mice with FUS-BBBO using a high peak negative pressure of 0.36 MPa against 8 specific brain regions within the striatum, thalamus, midbrain, and ventral hippocampus, which was within the range of pressures used for BBB opening in our previous studies^24, 28-30^. Lastly, we collected a blood sample after 7.5 min, which is the shortest time feasible for blood collection.

**Figure 2.**
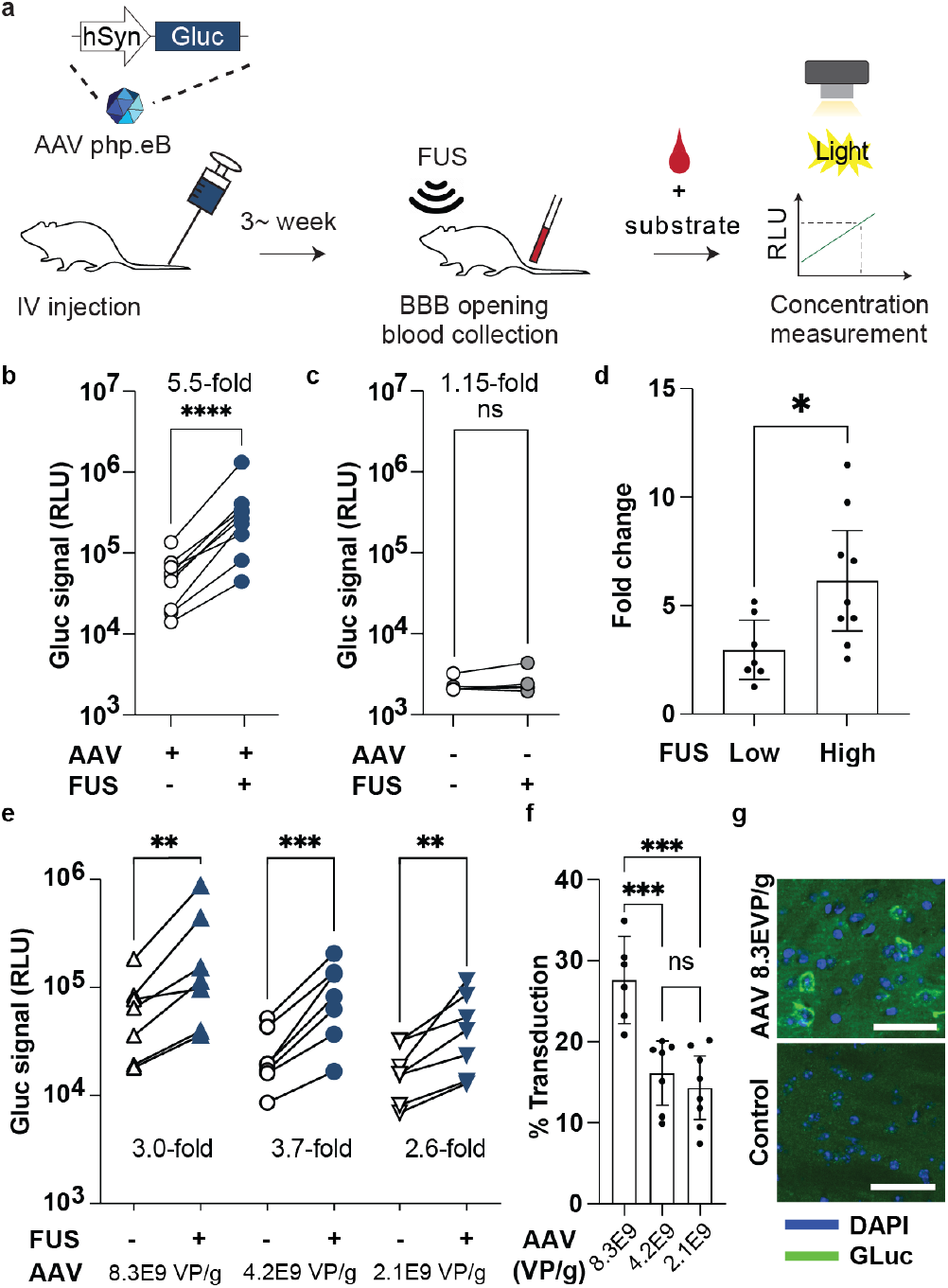
Recovery of GLuc expressed throughout the brain after administration of brain-tropic AAV. **(a)** Schematic for REMIS strategy, from i.v. injection of GLuc-AAV marker to bioluminescence readout of GLuc levels in the blood. **(b)** GLuc bioluminescence signal change before and after FUS-BBBO insonation in mice injected with GLuc-AAV (p<0.0001; two-tailed, ratio paired t-test; N=9). **(c)** Non-transduced mice were used as a control. (*P*=0.26; two-tailed ratio paired t-test, N=5). Serum RLU were measured by integrating bioluminescence signal from serum for 40 sec. **(d)** Foldchange in GLuc serum luminescence before and after FUS insonation with 0.36 MPa peak negative pressure (PNP) (high) or optimized PNP pressure ranging from 0.27 MPa to 0.33 MPa (low) depending on the targeted areas of the brain (P=0.023, two-tailed, unpaired t-test). **(e)** FUS-BBBO-induced release of GLuc from the brain for 2.1 × 10^9^ (*P*=0.0044; ratio paired t-test; N=7), 4.2 × 10^9^ (*P*=0.0002; ratio-paired t-test; N=7), and 8.3 × 10^9^ (*P*=0.0023, ratio paired t-test; N=7) viral particles per g of body weight using optimized peak negative pressures (ranging from 0.27 MPa to 0.33 MPa). **(f)** Comparison of transduction efficiency as measured by brain histology at the three AAV doses (*P*<0.001, one-way ANOVA, n=8, 7, 6, for low, medium, and high AAV dose groups respectively; Tukey HSD post-hoc test was used for pairwise comparisons; High dose vs medium: P=0.0009; High dose vs low: P=0.0002; Medium dose vs low: P=0.743). **(g)** Representative sections showing subcellular distribution of GLuc immunostaining (green) and nuclear stain DAPI (blue). Scale bar is 50 microns. ns > 0.05, *, *P*<0.05, **, *P*<0.01, ***, *P*<0.001, ****, *P*<0.0001.

Our results showed that all GLuc-AAV+ mice had increased GLuc signals in the plasma after insonation, with a mean fold change of 6.1±2.1 (95% confidence interval (CI); *P*< 0.0001, ratio paired t-test; N=9) (**Fig. 2b**). In the non-transduced control, we observed no changes in GLuc serum levels (*P*=0.26, ratio paired t-test, N=5).

To confirm BBB opening after insonation, we performed Evans Blue Dye (EBD) extravasation in the GLuc-AAV+ mice. On average, 7.3±0.4 out of 8 targeted brain sites in each mouse showed successful EBD delivery (95% CI; 92% of successful opening 66/72 of the targeted sites, N=9) (**Supplementary Fig. 1**). These regions represented approximately 4% of the brain volume in total. Taken together, our results thus far suggest that GLuc is successfully transported from the brain into the blood after insonation.

Next, although FUS-induced tissue damage typically self-resolves within hours to days^31^ and results in no neuronal loss^29^, we sought to minimize the potential for tissue damage while maintaining successful BBB opening and marker release. For this experiment, we reduced peak negative pressure levels down from 0.36 MPa by steps of 0.03 MPa and assessed the maximum radius of histological damage in different brain regions (**Supplementary Fig. 2a**). We defined damage as the presence of pockets of red blood cell (RBC) extravasation that typically observed after FUS-BBBO^32^. We graded this damage based on the size of the bounding ellipse within which all of the damage can be found, and defined 5 classes of damage – no damage (0 μm), or damages below 100 μm, between 100-200 mm, and then either between 200 and 400 μm, or greater than 400 μm. Typically, only a fraction of areas within that bounding box showed any discernible damage, both in our current study (**Supplementary Fig. 2b**) and previously-published work^32^. We chose the peak negative pressure levels to be 0.33 MPa in the striatum, 0.30 MPa in the thalamus and midbrain, and 0.27 MPa in the ventral hippocampus to enable successful opening in at least 7 out of 8 targeted sites in each mouse, while minimizing the presence of damage in the highest damage category (**Supplementary Fig. 2c**). We decided to use the lowest tested pressure for thalamus, where damage was present regardless of the tested pressure. Unless otherwise noted, we used these pressure levels for subsequent experiments.

We then investigated marker release before and after insonation under the optimized pressure levels and compared the marker release with 0.36 MPa (high) for each brain region. We found that lower optimized pressures still released GLuc, albeit at a lower fold-change over the baseline compared to high pressure (**Fig. 2d**, 3.0 vs 6.1 arithmetic mean fold-change for low and high pressure, *P*=0.0023, unpaired two-tailed t-test). However, lower pressure still resulted in highly significant release of the REMIS markers from the brain (**Fig. 2e**, left most panel, *P*=0.0002, N=7, ratio-paired t-test).

Afterwards, to evaluate whether transgene transduction efficiency affects GLuc signal levels in the serum post-insonation, we injected mice with GLuc-AAV at different doses (8.3 × 10^9^, 4.2 × 10^9^, and 2.1 × 10^9^ viral particles per g of body weight). After 3-4 weeks of expression, we tested GLuc serum levels before and after insonation. We found that, for each dosage group, all mice exhibited enhanced GLuc serum levels after insonation (**Fig. 2e)**. To measure the significance of the ratio of change before and after insonation we used a ratio paired t-test. The fold-changes compared to baseline were significant for each experiment injected with low, medium, or high dose AAV: 2.6fold ±1.3 (*P*=0.0044; N=7), 3.7-fold ±1.2 (*P*=0.0002, N=7), and 3.0-fold ±1.1 (*P*=0.0023; N=7) (mean, 95% CI; ratio paired ttest), respectively. Moreover, the ratio of post-insonation GLuc levels to baseline levels was unaffected by viral dose (**Supplementary Fig. 3**, *P*=0.4184; *F*=0.9148; one-way ANOVA). Further, histologic measurements of brain sections showed that high dose AAV mice resulted in 27.6% GLuc-positive cells, which was significantly higher than values observed with either low and medium viral doses (14.3% and 16.1%, P=0.0009, and P=0.0002, respectively) (**Fig. 2f**, *P*=0.0001; *F*=15.45; one-way ANOVA, pairwise comparisons P<0.001 when comparing high dose to lower doses, or P>0.05 otherwise.). Representative images showing transduction of Gluc are shown in **Fig. 2g**. These results suggest that ultrasound pressure, rather than the transduction efficiency, determines the efficiency of transgene product release from the brain.

Lastly, to confirm cell-type specificity of REMIS, we performed co-immunostaining for the expression of Gluc marker and the neuronal marker (NeuN). We found that 98.2% ±1.1 (95% CI) of Gluc-transduced cells were neurons (**Fig. 3a**). Representative images along with the examples of cells counted as NeuNare shown in **Fig. 3b** with arrowheads.

**Figure 3.**
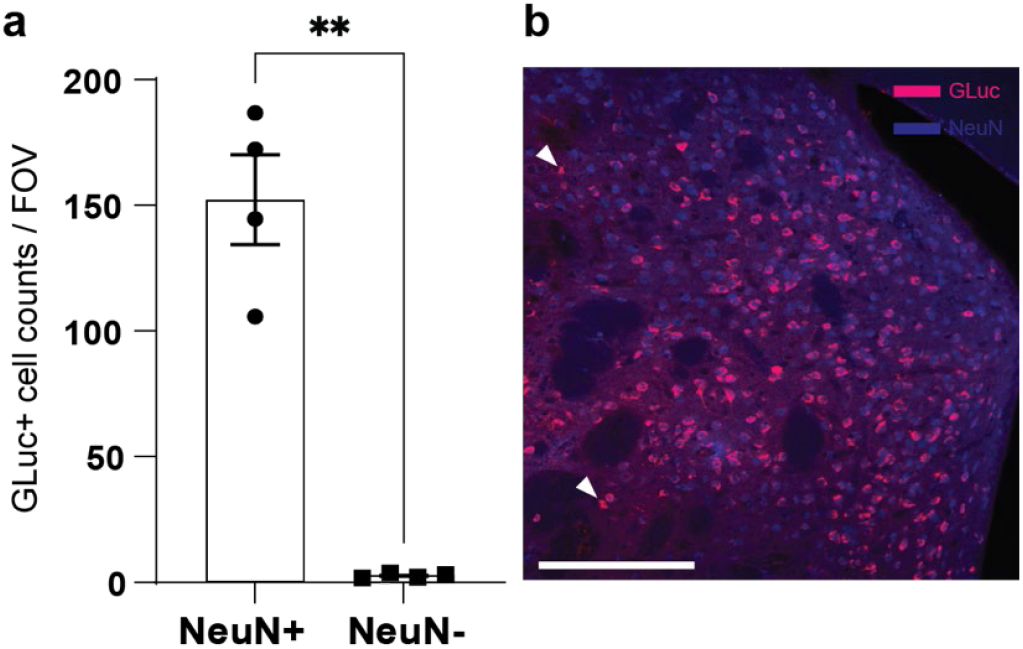
Cell-type specificity of REMIS. **(a)** We co-immunostained Gluc and the neuronal marker (NeuN) to quantify the number of cells expressing Gluc that are neurons and found that almost all were (98.2% ±1.1%, 95% CI) (paired t-test, p=0.0037). **(b)** Representative image of the brain section showing Gluc (magenta) and NeuN immunostaining (blue). Two of the cells positive in Gluc (Gluc+) and negative in NeuN (NeuN-) are designated with an arrowhead. Scale bar is 200 microns. ns > 0.05, *, *P*<0.05, **, *P*<0.01, ***, *P*<0.001, ****, *P*<0.0001.

### Released marker recovery pharmacokinetics after insonation

To evaluate the pharmacokinetics of marker released from the brain, we collected blood samples before and after insonation from GLuc-AAV+ mice (**Fig. 4a**). GLuc signal levels showed no statistically significant difference between 7.5 min and 120 min after insonation (**Fig. 4b**). This result could be explained by several possibilities, including the persistent release of markers from the brain and a marker half-life in the blood that is long compared to the duration of marker release from the brain.

**Figure 4.**
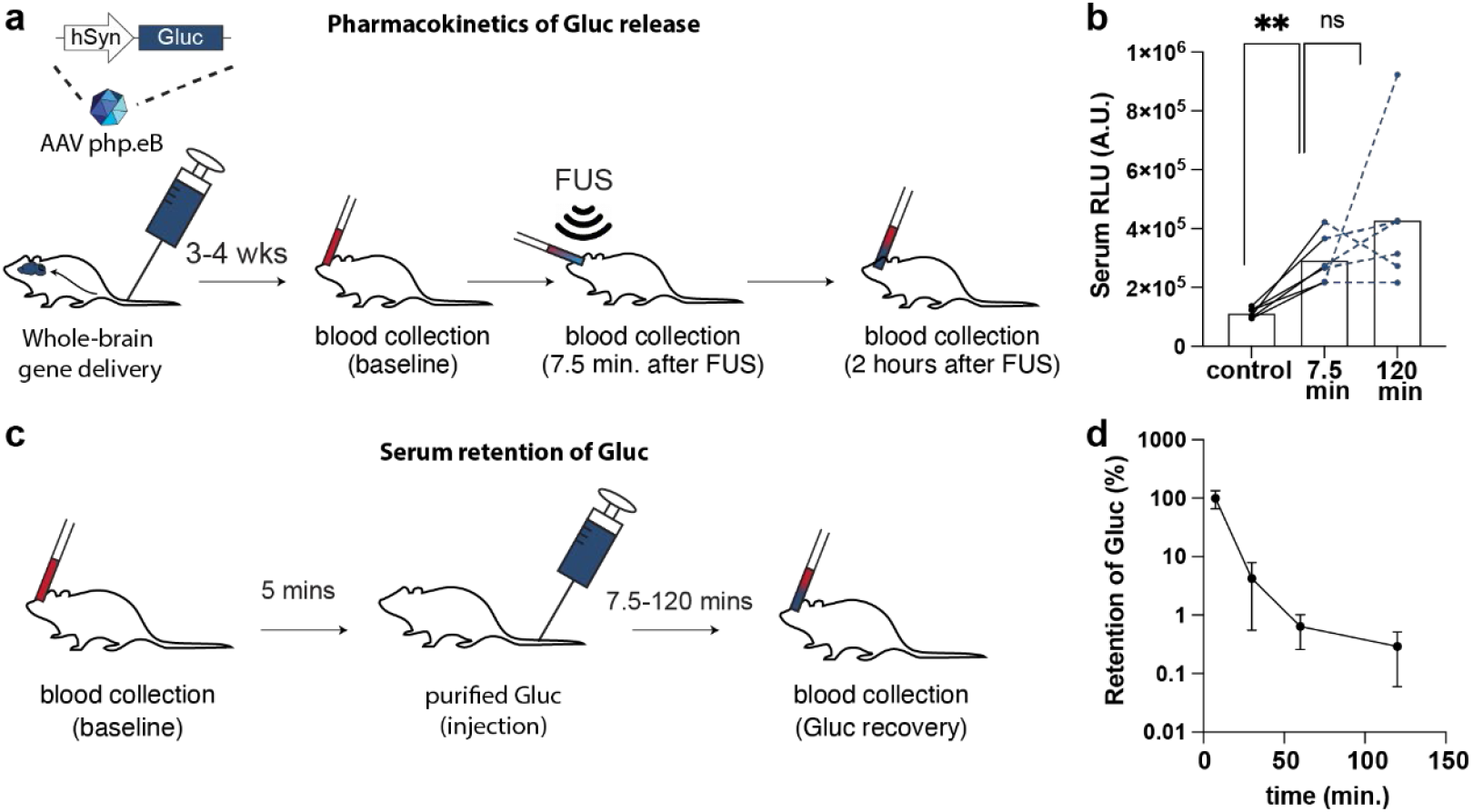
Pharmacokinetics of marker release in REMIS. **(a, b)** GLuc bioluminescence level measured at pre-insonation baseline, 7.5 min and 120 min after FUS-BBBO insonation (ns; ratio paired t-test; N=5). **(c, d)** Purified GLuc protein was administered i.v. to measure *in vivo* half-life of the protein in the blood. Blood was collected before injection as the baseline and then again at 7.5, 30, 60, and 120 min after injection to calculate the half-life (N=30 blood collections across 15 mice). ns > 0.05, *, *P*<0.05, **, *P*<0.01, ***, *P*<0.001, ****, *P*<0.0001.

To explore the possibility of a long marker half-life in the blood, we i.v. injected purified GLuc protein into mice and collected blood at different time points from 7.5 to 120 min (**Fig 4c**). We found that the GLuc half-life from a single exponential decay was 7.6±2.4 min (95% CI; N=30 blood collections in N=15 mice) (**Fig. 4d**), with over 99.4% of the GLuc eliminated by 60 min after injection. Rather than a long half-life, this result suggests a continuous release of GLuc from the brain over time, with serum levels of GLuc replenishing over 120 min, enabling a broad time window for blood collection for this marker.

### Noninvasive measurement of neuronal activity in specific brain regions

We next sought to determine whether REMIS could be used to measure endogenous neuronal activity in the brain. Here, we constructed a conditional genetic circuit to tether GLuc expression to neuronal activity (**Fig. 5a**). To gain temporal control over Gluc, we used designer receptor exclusively activated by designer drug (DREADD) hM3Dq, which induces neuronal firing when turned on by the DREADD agonist clozapine-*N*-oxide (CNO)^7^. We also incorporated into the circuit the doxycycline (Dox)-dependent Tet-Off system called Rapid Activity Marking (RAM)^33^, which requires both an active *c-Fos* responsive promoter and the absence of Dox to drive Gluc expression that is then released into the interstitial space of the brain and a nuclearly-localized GFP under internal ribosome entry site (IRES) as a histological ground-truth control of *c-Fos* activation (**Fig. 5b**).

**Figure 5.**
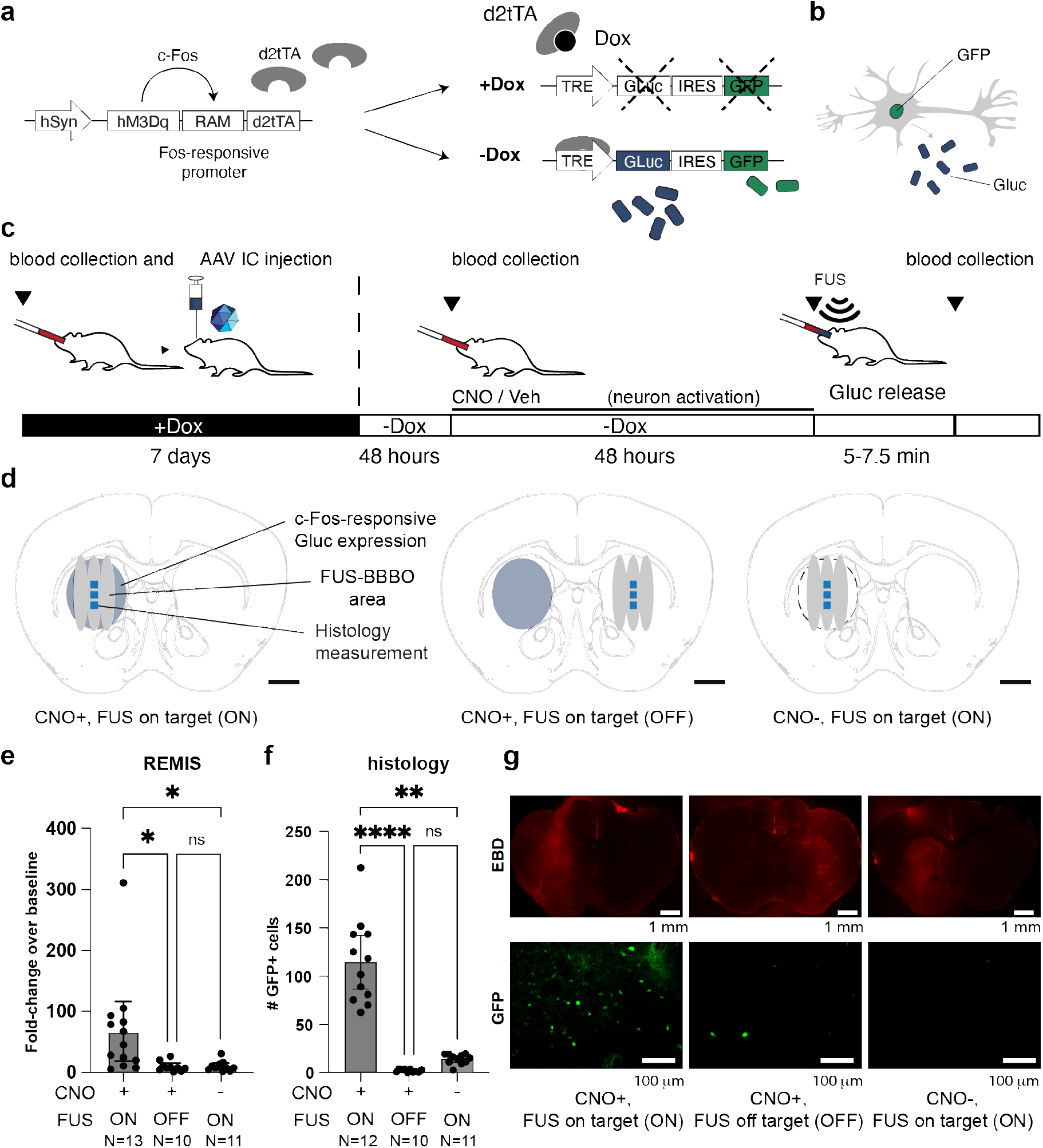
Noninvasive measurement of c-Fos specific brain regions with REMIS. **(a)** Schematic of the experimental procedure using doxycycline (Dox)-dependent Tet-Off Rapid Activity Marking (RAM). GLuc was placed under control of RAM to express GLuc following prolonged neuronal activity. As an independent control for measuring c-Fos GFP was also expressed under Internal Ribosome Entry Sites (IRES). To activate the neurons expressing RAM and GLuc, an activatory DREADD, hM3Dq, was placed in cis with the RAM system’s promoter. To ensure temporal precision of c-Fos recording, RAM system can be shut down with Doxycycline (Dox) or begin recording when Dox is withdrawn. **(b)** Upon neuronal activation, Gluc is expressed and released into the interstitial space of the brain, while GFP is expressed intracellularly and localized to the nucleus of neurons. **(c)** Blood was drawn as a baseline and followed by an injection of AAV carrying GLuc expressed under RAM system into unilateral side of caudate putamen in mice. Dox was administered for 7 days in food to allow for expression of RAM system components, while preventing *c-Fos*-responsive production of GLuc. Dox food was then withdrawn and, 48 h later, blood was collected and either a chemogenetic prodrug Clozapine-N-Oxide (CNO) or a vehicle control was injected intraperitoneally (i.p.). After 48 h, FUS-BBBO was performed to release GLuc into the blood and blood was drawn. **(d)** Illustration of on- and off-target FUS-BBBO sites in the striatum for each group in gray. *C-Fos* expression is shown in blue. Histology measurements were performed within three regions located within the opened areas and shown in dark blue. **(e)** GLuc levels in the serum after opening the BBB of target regions (*P*=0.0032; Kruskal-Wallis test, Kruskal-Wallis statistic=11.48, with multiple comparisons; CNO + FUS on target vs CNO + FUS off target: P=0.0104; CNO-on target vs Vehicle, P=0.0137; CNO + FUS off target vs Vehicle: P>0.9999). **(f)** Ground-truth histology measurement of GFP expressed under IRES in response to c-Fos accumulation. (*P*=0.0001; Kruskal-Wallis test, Kruskal-Wallis statistic=27.55, with multiple comparions; CNO + FUS on target vs CNO + FUS off target: P<0.0001; CNO-on target vs Vehicle, P=0.0099; CNO + FUS off target vs Vehicle: P=0.0636). **(g)** Representative images showing the EBD signal (top) or the GFP expression in the tissue sections (bottom). ns > 0.05, *, *P*<0.05, **, *P*<0.01, ***, *P*<0.001, ****, *P*<0.0001.

To activate neurons, we injected AAVs carrying the entire *Fos*-responsive RAM-controlled Gluc expression system into the left caudate putamen of mice. We then administered Dox (P.O. in food *ad libitum*) for 7 d to suppress Gluc expression until a specified window needed for neuronal activity recording. Then, 48 h after withdrawing Dox, we collected a baseline blood sample (**Fig. 5c**). Immediately following, we injected CNO at a previously-validated dose^34^ intraperitoneally (i.p.) to drive neuronal activation and the coupled expression of Gluc. CNO, a prodrug^7, 35^ for the DREADD activator in vivo, was chosen over other chemogenetic activators due to its long timeline action^34^, well-suited to long-term recording with the RAM system. As a control, we also injected a group of mice with vehicle to evaluate baseline levels of GLuc signal in the absence of CNO-induced neuronal activity. After 48 h^19, 23^, we insonated mice at a target site within the striatum (N=10 for vehicle and N=13 for CNO+ mice) (**Fig. 5d, left and middle)**. To test the site-specificity of REMIS, we also insonated a group of CNO+ mice in the contralateral striatum (N=11) (**Fig. 5d, right**).

Following on-target insonation, we found that DREADD-activated neurons in CNO+ mice showed 67.6-fold ±44 (mean with 95% CI) increase of GLuc in the blood. This enrichment was higher than that observed in both the vehicle group (8.3-fold ±3.1, mean with 95% CI, *P*=0.0137, Kruskal-Wallis test, with multiple comparisons) and off-target CNO+ group (8.1-fold±6.3, mean with 95% CI, *P*=0.0104, Kruskal-Wallis test, with multiple comparisons) (**Fig. 5e)**. Moreover, targeting the contralateral site was not significantly different from targeting the striatum in the vehicle control (*P*>0.9999, Kruskal-Wallis test), showing site-specificity of REMIS.

We then examined whether measurements of neuronal activation using REMIS (**Fig. 5e**) could be recapitulated histologically. The *c-Fos*-responsive promoter^23^ of our genetic circuit is designed to drive concurrent expression of secreted GLuc and intracellular green fluorescent protein (GFP) in response to induced neuronal activity (**Fig. 5a, b**). Our data showed that CNO+ mice that expressed GLuc in the striatum had the highest number of GFP-positive cells within the insonated target areas when compared to the off-target mice and vehicle controls (**Fig. 5f**). Specifically, we found an 8.0-fold difference in the number of GFP-positive cells between on-target CNO+ mice and the vehicle control (*P*<0.0001, Kruskal-Wallis test), which corresponds to the 8.1-fold difference between these two groups when GLuc was measured using REMIS (**Fig. 5e)**. Representative images highlighting the area of BBB opening and GFP expression for each group is shown in **Fig. 5**. These results suggest that REMIS can measure *c-Fos* activation in specific brain regions.

## DISCUSSION

Taken together, our results establish REMIS as a paradigm for noninvasive, site-specific measurements of transgene expression in genetically-targeted cells. Instead of directly measuring gene expression within the deep tissue, which is difficult to achieve, REMIS provides a means of recording reporter concentration in an ultrasound-defined region by transporting engineered markers of gene expression into the blood where they can be easily quantified. Our results demonstrate the suitability of this approach for the confirmation of gene delivery and investigating cellular physiology, specifically for monitoring sustained neuronal activation levels that give rise to *c-Fos* activity.

Compared to existing approaches, REMIS has several advantages. Traditionally, brain biopsy or post-mortem histology is used to extract cells from the brain and measure gene expression. However, they are invasive and destructive to the tissue. In consequence, they cannot record gene expression activity over multiple timepoints and they damage the very brain region being studied. REMIS represents a nondestructive and potentially repeatable^36^ alternative to biopsy-based measurement of gene expression.

Second, whereas imaging methods such as MRI with genetically-encoded contrast agents can visualize the entire brain, REMIS enables monitoring of a pre-defined brain region with millimeter precision. REMIS, however, has two important advantages. First, it achieves over an order of magnitude higher signal to baseline ratios than observed for genetically-encoded MRI contrast agents^4, 5, 37,38^. Additionally, REMIS measures molecule concentration through biochemical blood testing, enabling inexpensive detection of multiple types of molecules, possibly in a single sample.

Compared to PET, REMIS does not rely on the development of transgene-binding radioactive BBB-permeable probes, while maintaining comparable spatial precision. When compared to optical imaging, REMIS allows measurement of markers in any brain region that is accessible with focused ultrasound, including behind thick skulls in various species or in deep brain regions^24, 36, 39, 40^ both locally or brain-wide^29, 41^. In contrast, for optical methods, such as fluorescence or bioluminescence imaging, depth of penetration and scattering of light through the skull or tissue are major technical obstacles. Thus, REMIS has both an advantage over optical methods and future utility in large animal species and deep or large brain regions.

Compared to RMAs^19^, REMIS appears less sensitive with lower fold changes over the baseline and requires ultrasound procedure before readout. In contrast, RMAs cross through an intact BBB on their own, allowing for a simpler readout. However, RMAs rely on the site-specific gene delivery to determine the spatial precision and averages signal over the entire transduced cell population. REMIS, on the other hand, can sample different brain sites within the transduced cell populations, providing it with unique advantages. For example, REMIS can be used for validating gene delivery sites, assessing a diffusion of secreted transgenes^42, 43^, or monitoring gene expression in individual brain sites. Current brain gene therapies struggle with monitoring gene expression without invasive procedures. REMIS offers a non-surgical, non-tissue-destructive solution by augmenting gene therapy to express detectable REMIS markers, akin to using fluorescent proteins in histology. REMIS also could enable long-term spatially-specific transgene expression monitoring, without causing tissue damage, as shown by safety of multiple applications of FUS-BBBO in large animals^36^.

Our results showing noninvasive monitoring of *c-Fos* in a specific brain region could also be applied to a range of studies. Immediate early genes (IEG) such as *c-Fos* and *Arc* are activated by heightened neuronal activity^44^ and, thus, REMIS could be used to point to a brain region of interest and measure long-term changes in neuronal activity, for example to d etermine successful neuromodulation, or activation neuronal ensembles associated with learning^45^ and memories^46^.

The spatial resolution of REMIS is dependent upon the parameters of focused ultrasound for the BBB opening and ranges from millimeter to sub-millimeter precision^24, 40^. In humans, current devices open the BBB with millimeter precision^40^.

One possibility that could reduce the resolution is diffusion of the markers away from the site of their expression. Our experiments suggest that the diffusion is limited in the tested scenario. We measured *c-Fos* activation in the striatum contralateral to the area of activation (distance of ~2.5 mm) and found no diffusion of markers into that region. Moreover, the application of FUS-BBBO to the striatum contralateral to the CNO-activated brain region had an effect that was indistinguishable from opening the BBB in vehicle control mice (**Fig. 5e**).

REMIS’s cell-type specificity is reliant on the genetic targeting. In our results, we used synapsin promoter which restricts expression of the REMIS markers to neurons, and found their expression was highly specific (**Fig. 3**). The fold-change of REMIS markers in the serum is dependent on the ultrasound pressure (**Fig. 2d**), rather than the viral dose (**Supplementary Fig. 3**). The process of marker recovery was robust – of note, every single mouse tested in this study has shown an increase in the REMIS marker levels after the BBB opening, reaching high levels of significance even for lower fold-changes (**Fig. 2e**). These results suggest that REMIS could be successfully used in small cohorts of animals, which is critical for large animal studies, or any potential clinical applications.

Finally, the temporal resolution of REMIS is largely dependent on the temporal profile of gene expression driving it. The release of markers from the brain itself is rapid with markers appearing in serum within minutes (**Fig. 4a, b**). The c-Fos^34^ expression, on the other hand, occurs over minutes to hours timescale, with RAM system needing 24-48 hours for detectable expression^33^. Importantly for the convenience of measurement we saw a steady, continuous, release of the GLuc over at least 120 minutes. Steady release makes the experimentation less time-dependent and thus more robust. Other markers could be designed that exit the brain more rapidly depending for example on their size and diffusivity, or binding affinity within the extracellular matrix of the brain.

The REMIS system could be made more powerful with improvements to each of its components: FUS-BBBO, methods of gene-delivery to the brain, gene circuits measuring endogenous gene expression activity, or methods of serum marker detection. For example, real-time monitoring of cavitation^47-50^ has been widely used in human FUS-BBBO^51^ studies to optimize the ultrasound pressures for safety and account for differences in skull attenuation. Design of new AAV vectors can lead to improved noninvasive delivery. For example, AAVs can be designed to pass through an intact BBB after a systemic injection and specifically transduce the central nervous system^52^, or be more efficiently and tissue-specifically delivered with FUSBBBO^53^. They can also be optimized to work in different species^54, 55^. These noninvasive gene delivery tools synergize with REMIS which enables similarly noninvasive measurement of the success of gene delivery. Finally, design of new gene circuits^33, 56, 57^, molecular recorders^58, 59^, or mRNA sensors^60, 61^, that can translate cellular physiology into gene expression will enable new applications of noninvasive techniques like REMIS. REMIS uses biochemical testing which opens up the possibility of measuring multiple types of markers simultaneously, potentially enabling multiplexed monitoring through detection techniques such as mass spectrometry proteomics^62^ or single-molecule protein sequencing^63, 64^. Such multiplexed imaging is not currently possible with MRI, PET, or even with optical methods, where the number of colors in *in vivo* imaging is orders of magnitude fewer than what can be detected through serum sampling with proteomic techniques.

These improvements will facilitate the development and translation of REMIS as a paradigm for precise non-invasive monitoring of genetically-targeted cell populations within specific brain regions.

## MATERIALS AND METHODS

### Animals

Wild-type C57BL/6J (Strain #000664) male and female mice were obtained from Jackson Laboratory (Bar Harbor, ME). Animals were housed in a 12 h light-dark cycle and were provided with food and water ad libitum. All experiments were conducted under a protocol approved by the Institutional Animal Care and Use Committee of Rice University.

### Plasmid construction

To construct AAV-hSyn-GLuc, the vector AAV-hSyn-GLucM23-iChloC-EYEP (Addgene #114102) was digested with SpeI and EcoRV (New England Biolabs) to isolate the backbone containing the hSyn promoter. GLucM23, a GLuc variant, was amplified by PCR from the same vector and its DNA was extracted using the Monarch DNA Gel Extraction Kit (New England Biolabs, Ipswich, MA). GLuc was inserted into the digested backbone through Gibson Assembly Hi-Fi kit (New England Biolabs, Ipswich, MA). AAV-hSyn-hM3Dq-RAM-d2tTA was constructed as previously described^19, 23^. To construct AAV-RAM-GLuc-IRES-GFP, the GLuc DNA was amplified from the AAV-TRE-RMA-IRES-GFP plasmid in our previous work^19, 23^ and assembled into the same backbone after digestion using PmeI and BamHI. AAV-hSyn-CLuc was constructed similarly by amplifying the CLuc DNA from CLuc-RMA (CLuc-Fc)^19, 23^ plasmid and inserted into the same backbone after digestion using EcoRV and BamHI.

### Adeno-associated virus production

PHP.eB^65^ AAV virus was packaged with the AAV-hSyn-GLuc plasmid construct using a commercial service (Vector Builder, Chicago, IL) and the titer was provided by the manufacturer.

### Intravenous administration of AAVs

AAV was injected intravenously (i.v.). Mice at 10–14-weeks old were anesthetized with 2% isoflurane in oxygen and then cannulated in the tail vein using a 30-gauge needle connected to PE10 tubing. The cannula was then flushed with 10 units (U) ml^-1^ of heparin (Sigma Aldrich) in sterile saline (Hospira) and affixed to the mouse tail using a tissue adhesive. Subsequently, the mice were injected via tail vein with PHP.eB AAV (2.1-8.3 × 10^9^ viral particles per g of body weight) encoding GLuc under the hSyn promoter. AAV injected mice were housed for 3-4 weeks to allow for gene expression.

### Focused Ultrasound BBB opening

Mice at 14–18-weeks old were anesthetized with 1-2% isoflurane in oxygen. The hair on mice head was removed by shaving with a trimmer. The mice were then cannulated in the tail. Subsequently, the mice were placed on a stereotaxic mount, with their heads held in place with a custom-made plastic nosecone and secured with ear bars. Bregma and Lambda markers were used to target ultrasound to specific brain regions using a stereotaxic frame (RK-50, FUS Instruments). Ultrasonic transducer was coupled to the shaved area of the head via degassed water in water bath and degassed aquasonic ultrasound gel. The mice were injected with approximately 2.8 × 10^6^ DEFINITY microbubbles (Lantheus) dissolved in sterile saline, per g of body weight for each targeted site. For each targeted site, DEFINITY was re-injected before insonation. The ultrasonic parameters used were 1.5MHz, 10 ms pulse train length, 1Hz pulse repetition frequency for 120 pulses. The pressure for insonation was varied between experiments and targeted sites based on evaluation of safety and efficacy of BBB opening in this study. The range of peak-negative-pressure used in this study was 0.27-0.36 MPa, as calibrated by the manufacturer (FUS Instruments, Toronto, Canada). In the GLuc recovery experiment, targeted sites represented 4% of the brain volume in total, as measured by full-width half-maximum pressure of the FUS field. Due to equipment failure, experiments on c-Fos measurement used a new transducer with a different calibration curve. BBB opening and tissue damage were used to confirm the validity of new calibration in vivo and used manufacturer-calibrated pressure of 0.46 MPa, which is not directly comparable with the pressures used in previous experiments due to differences in manucturer’s (FUS instruments, Toronto, Canada) calibration methods. However, the opening was present and comparable to previous experiments (**Fig. 5g**) with 91.2% (31/34) mice showing opening (**SI Fig. 4**) and we observed no apparent damage in N=12 randomly selected mice (N=4 per group for CNO with on target FUS-BBBO, CNO with off target FUS-BBBO, and vehicle with on-target FUS-BBBO). After ultrasound insonation, aquasonic ultrasound gel was removed and the incision was closed. Evans Blue Dye (5%) dissolved in 1X PBS was injected i.v. EBD passes selectively through a permeable BBB and localizes in the brain parenchyma, allowing visualization of where the BBB has been opened^66^. After 20 min, mice were perfused with 10% neutral buffered formalin (Sigma-Aldrich) after displacing blood with 10 units (U) ml^-1^ of heparin in 1X PBS via cardiac perfusion.

### Pharmacokinetic analysis of GLuc serum half-life

Purified GLuc protein (Nanolight Technology) was injected i.v. through a tail vein catheter. C57Bl6j mice at 10–14-weeks old were injected with purified GLuc protein in 1X TBS under isoflurane anesthesia (1-5% in oxygen). After a period of 7.5-120 minutes, mice were again anesthetized in 2% isoflurane in oxygen. Afterwards, 1-2 drops of 0.5% ophthalmic proparacaine were applied topically to the cornea of an eye and a heparin-coated microhematocrit capillary tube (Fisher Scientific) was placed into the medial canthus of the eye and the retro-orbital plexus was punctured to withdraw 50-100 μl of blood. The collected blood was centrifuged at 5,000g for 15 min to isolate plasma and stored at -20°C until use. To conduct luciferase assay, 15 μl of plasma was placed in a black 96-well plate. Bioluminescence of GLuc was measured by injecting plasma samples with 50 μl of 20 μM coelenterazine (CTZ) (Nanolight Technology) dissolved in luciferase assay buffer using an injector in a Tecan M200 microplate reader (Männedorf, Switzerland).

### Immunostaining

Mice brains were extracted and postfixed in 10% neutral buffered formalin overnight at 4°C. Coronal sections were cut at a thickness of 50 μm using a vibratome (Leica) and stored at 4°C in 1X PBS. Sections were stained as follows: 1) pre-incubate with permeabilizing agent (1% Triton X-100, 0.5% Tween 20 in 1X PBS) for 1 h; 2) block for 1 h at room temperature with blocking buffer (0.3% Triton X-100 and 10% normal donkey serum in 1X PBS); 3) incubate with primary antibody overnight at 4°C; 4) wash in 1X PBS for 10 min 3 times; and 5) incubate with secondary antibody for 4 h at room temperature. After last washes in 1X PBS, sections were mounted on glass slides using the mounting medium (Vector Laboratories) with or without DAPI and cured overnight in dark at room temperature. Antibodies and dilutions used are as follows: rabbit anti-GLuc (1:1,500, Nanolight Technology) and Alexa 488 secondary antibody (1:500, Life Technologies). All images were acquired by the BZ-X810 fluorescence microscope (Keyence).

### Statistical analysis

Two-tailed paired t-test with unequal variance was used to compare two data sets when comparing means, or ratio-paired t-test was used to compare fold changes between two groups. Oneway ANOVA or Kruskal-Wallis tests with Tukey honestly significant difference post hoc test was used to compare means between more than two data sets, depending on whether the standard deviations of compared groups were significantly different in Brown-Forsythe test. Kruskal-Wallis was selected for c-Fos measurement data due to showing significantly non-normal distribution (Shapiro-Wilk test for CNO-on-target FUS group, P=0.0006). All *P* values were determined using Prism (GraphPad Software), with the statistical significance represented as ns (not significant), \**P* < 0.05, \*\**P* < 0.01, \*\*\**P* < 0.001, \*\*\*\**P* < 0.0001. The sample and effect sizes were calculated using G*power 3 software^67^, assuming 80% power, and alpha value of 0.05. In the absence of previous data that could be used for calculation of effect size we relied on sample sizes used in invasive measurement of tissue gene expression^68^ (for data in Fig. 2a, Fig. 3), or in noninvasive chemogenetic neuromodulation with gene delivery using FUS-BBBO^24^ (for data in Fig. 5e, fg).

### Stereotaxic injection of AAV

AAV were injected into the brains using a microliter syringe equipped with a 34-gauge beveled needle (Hamilton) installed to a motorized pump (World Precision Instruments) using a stereotaxic frame (Kopf). For AAV, PHP.eB serotype was used. For chemogenetic neuromodulation experiments, three different AAVs were delivered. AAV encoding hSyn-hM3Dq-RAM-d2tTA and RAM-GLuc-IRES-GFP were delivered to the left hemisphere caudate putamen (CPu) in the striatum in the following ipsilateral coordinates: (AP +1.0 mm, ML +2.0, DV +3.8 mm), (AP +0.5 mm, ML +1.5, DV +3.0 mm), (AP -0.5 mm, ML +2.5, DV +3.5 mm). AAV encoding hSyn-CLuc was delivered to the right hemisphere CPu in the striatum in the contralateral coordinates. For each mouse, 1.0 × 10^9^ viral particles of each AAVs were delivered correspondingly. For AAV injections, 568 nl AAV (hSyn-hM3Dq-RAM-d2tTA) and 116 nl AAV (RAM-GLuc-IRES-GFP) were prepared in a cocktail and 684 nl total volume was injected each site. AAV (hSyn-CLuc) alone was injected 500 nl each site. The injection speed was 250 nl min^-1^. The needle was kept at the injection site for 10 min owing to the relatively high volume of injection.

### Doxycycline control of gene expression

For chemogenetic neuromodulation experiments, mice were placed on 40 mg kg^-1^ of Dox chow (Bio-Serv) 24 h prior to intracranial injection of AAVs. Dox chow was maintained for 7 d after the AAV injection. Dox chow was removed 48 h prior to inducing chemogenetic activation.

### Chemogenetic activation of neuronal activity

Water-soluble CNO (Hello Bio #HB6149) was dissolved in saline (Hospira) at 1mg ml^-1^ and stored at -20°C until use. To induce chemogenetic activation of mice expressing hM3Dq, CNO was injected intraperitoneally (i.p.) at 5 μg per g mouse, respectively.

### Blood collection for luciferase assay

For the groups of mice in GLuc *in vivo* half-life measurement experiment, baselines were collected 5 min prior to GLuc i.v. injection. Experimental samples were collected 7.5 min post-injection in common and 30, 60, and 120 min post-injection based on their assigned groups. For the groups of mice measuring transduction after systemic AAV delivery, baselines were collected before subcutaneous analgesic drug injection in the ultrasound insonation procedure. Experimental samples were collected 7.5 min after an ultrasound insonation session. For the group of mice in GLuc recovery in extended time frame experiments, 120 min timed experimental samples were collected 7.5 min or 120 min after an ultrasound session. For the groups of mice where we used REMIS to measure chemogenetic activation, baselines were collected twice; before intracranial AAV injection and before CNO i.p. administration. Experimental samples were collected 7.5 min after an ultrasound insonation session.

### Hematoxylin staining

Sectioned brain samples were stored in 1X PBS and prepared in water 10 min prior to staining. Samples were dehydrated in 100% ethanol and 95% ethanol, 1 min each. Then, the samples were rinsed in water for 1 min. Next, samples were immersed in 0.4x diluted Mayer’s Modified Hematoxylin solution(Abcam, #64795) for 1 min and rinsed again in water for 3 min. Nuclear stain was completed by immersing samples in the Blueing reagent(Abcam, #66152) for 20 sec. Finally, samples were rinsed in water for another 3 min. Samples were rehydrated in water for another 10 min before mounting.

### Cell positivity quantification

*C-Fos* images were taken on a Keyence BZ-x810 using the 20x objective lens. Each brain section had three images taken, one on top of the other in a vertical line inside of the blood brain barrier opening guided by the EBD signal (**Fig. 5d**). Images were counted manually in Zen Software (J.S.T.). Each image was loaded in and the upper limit of the histogram was reduced to 15,000 out 65536 for every image to aid in visualization of all cells. Blinding was not possible due to the stark differences between the experimental group and controls.

Gluc cell counts images were taken on a Keyence BZX810 using the 20x objective lens. Single hemisphere images were taken and counted in zen software by adding a horizontal only grid line spaced by (400 pixels) vertically between each line. Only green and blue cells that touched this line were counted. Green cells were taken in the 488 channel and designate Gluc expression. Blue cells were taken in the Dapi channel and were stained using DAPI containing mounting media. The experimenter (R.W.) was blinded to identity of the groups.

Cell specificity for Gluc and NeuN positive images were taken on a Zeiss LSM-800 Confocal microscope. NeuN was stained using Novus Biologicals RBFOX3/NeuN antibody conjugated to Alexa Fluor 405. GLuc stained with anti-GLuc antibody (nanolight technology) and an Alexa Fluor 594 secondary antibody. GLuc and NeuN images were taken using the 20x objective lens on the confocal, each image was overlaid and counted manually using zen software. Total of 11 sections across 4 mice was stained and counted (2-3 sections per mouse).

## ACKNOWLEDGEMENTS

This work was partially supported by the Michael J. Fox Foundation, grant #020154 to J.O.S., and NIH grant R21EB033059 to J.O.S, and the Welch Foundation grant # C-2048-20200401 to J.O.S. Related work in the lab was performed under Packard Fellowship for Science and Engineering to J.O.S (#2021-73005).

## AUTHOR CONTRIBUTIONS

J.O.S. and J.P.S. conceived and planned the research. J.P.S, J.S.T, S.L., and Z.H. performed the in vivo experiments. J.S.T, and J.P.S, performed the histological experiments, S.L. helped design and implement c-Fos measurement experiments, J.O.S, J.P.S, and J.S.T., R.Z.W. analysed the data. J.O.S. and J.P.S. wrote the manuscript with input from all other authors. J.O.S. supervised the research.

## Competing interests

The authors declare no competing financial interests.

## Notes

### Competing Interest Statement

The authors have declared no competing interest.

