## Supplementary figures for "Acoustically-Targeted Measurement of Transgene Expression in the Brain"

### Supplementary data

### Supplementary Figures

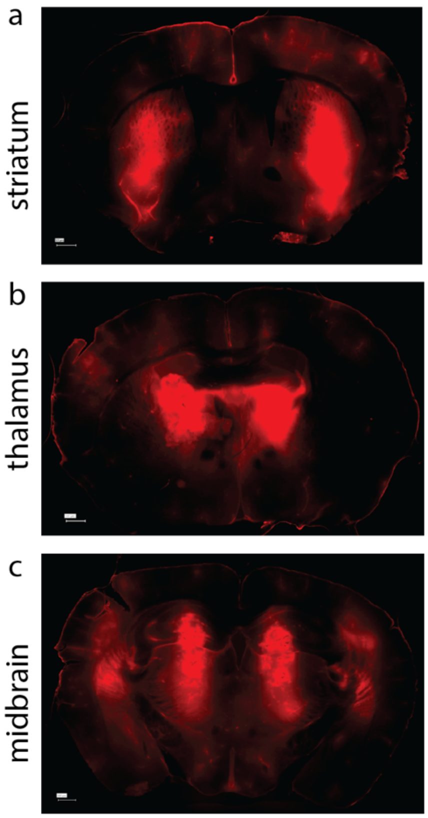


**Supplementary Figure 1. BBB opening in high ultrasound pressure group AAV(+) FUS(+)**. Representative brain section on red channel visualizing EBD extravasation and approximating the targeted regions of the brain. The volume targeted was calculated from the full-width-half-maximum (FWHM) pressure profile, which was approximated as an an ovoid with the length of 5.4 mm and width of 0.9 mm, for a total volume of 2.3 $mm^3$. Assuming the C57BL6j mouse brain volume of 508.91 $mm^3$ [69], 8 sites at 2.3 $mm^3$ each would result in 3.6% of the targeted volume. Scale bars are 500 microns.

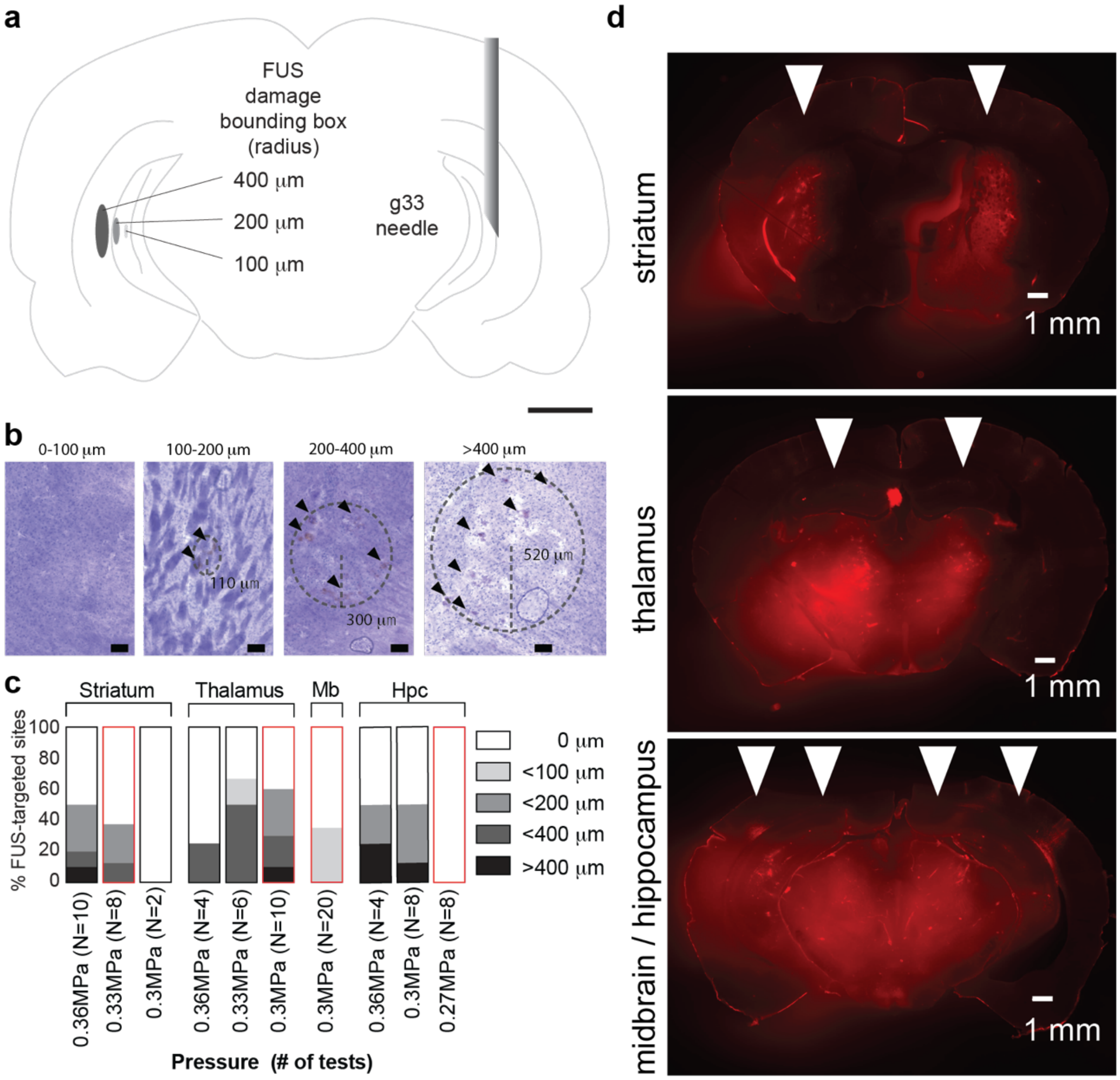


**Supplementary Figure 2. Optimization of FUS peak negative pressure levels for different brain regions**. (a) The size of areas within which any damage is contained are shown for illustration purposes and have the approximate shape of an ultrasound pressure field. Throughout this area, only some spots show damage. For comparison, a g33 needle is shown to visualize the tissue displacement needed for molecule delivery or biopsy. (b) Quantification of the maximum radius of the histological damage observed in mice. Histological damage was measured in 4 different areas of the brain; striatum, thalamus, midbrain (Mb), and hippocampus (Hpc). Optimal peak negative pressures were adjusted by steps of 0.03MPa down from 0.36MPa to find a set of conditions that resulted in no or minimal damage while maintaining BBB opening. Conditions shown in red boxes were chosen for all experiments, unless otherwise noted. **(d)** Representative images showing extent of the BBB opening (EBD extravasation, red, highlighted with arrowheads) with the chosen parameters. Scale bars are 1 mm.

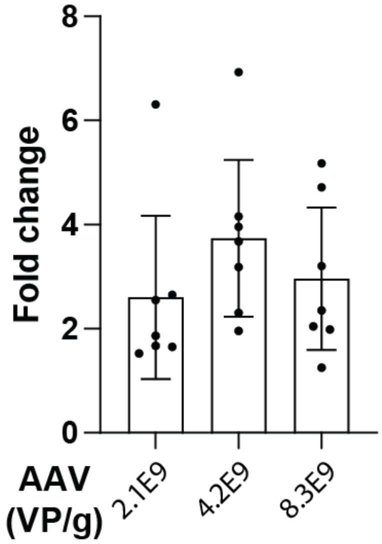


**Supplementary Figure 3. Comparison of fold changes in serum concentration of GLuc induced by FUS-BBBO**. AAV php.eB carrying GLuc under hSyn1 promoter was injected intravenously at different doses 2.1 x $10^9$ viral particles (VP) per gram of body weight, or double (4.2x$10^9$ VP/g) or quadruple that dose (8.4x$10^9$ VP/g). After 3 weeks, FUS-BBBO was performed to release GLuc from the brain. The pressure used for release was optimized for safety, and included 0.33 MPa in the striatum, 0.3 MPa in the thalamus and midbrain, and 0.27 MPa in the hippocampus ($P$=0.4184; $F$=0.9148; one-way ANOVA).

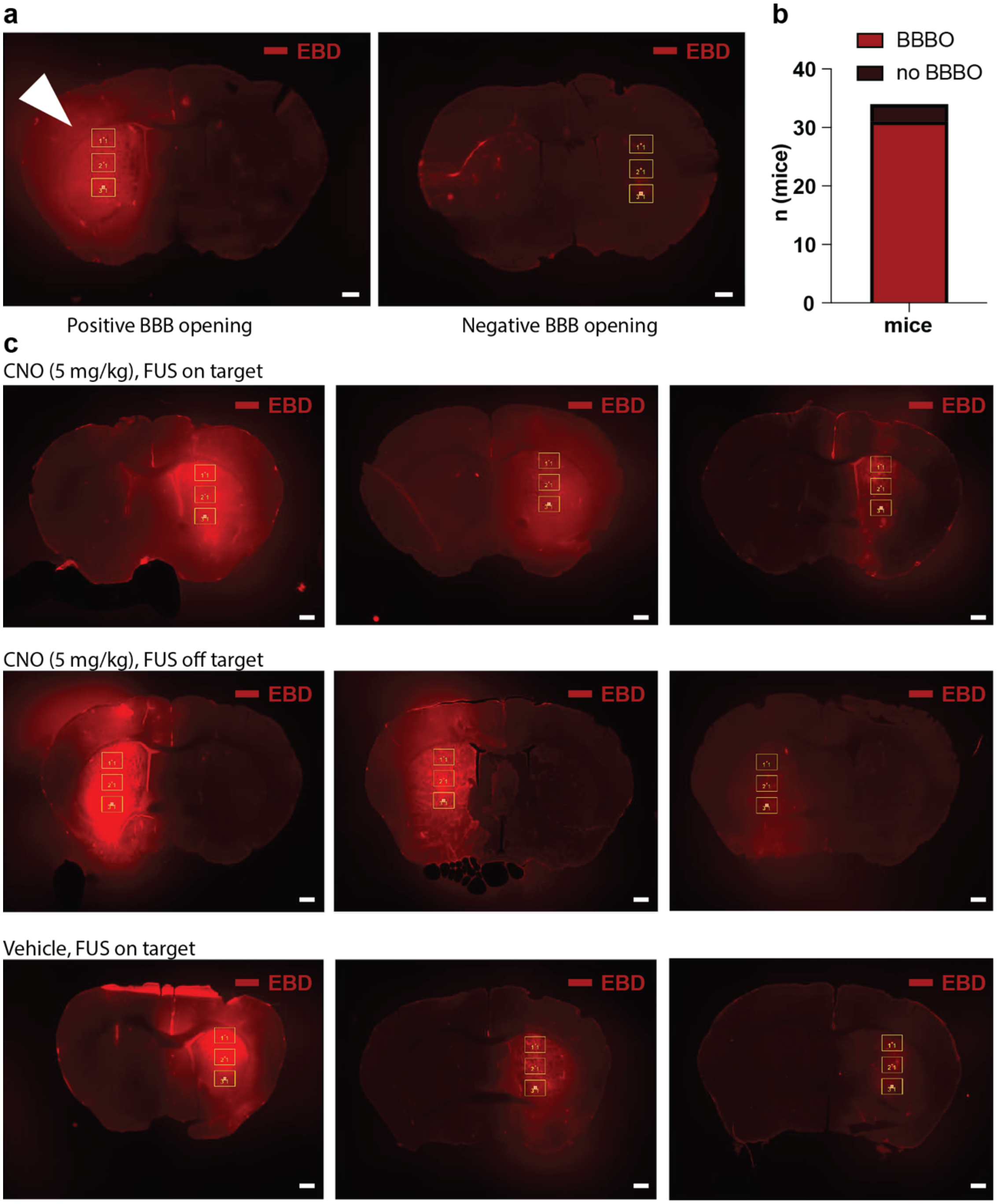


**Supplementary Figure 4. Validation of the BBB opening during measurement of *c-Fos* activity with REMIS. (a)** Within 20 minutes of the BBB opening, mice were adminterestered evans blue dye (EBD, red) I.V. which can cross through an opened BBB from the blood into the brain. EBD shows red fluorescence that can be assessed to evaluate the quality of BBB opening. The presence or lack of BBB opening was assessed qualitatively as positive or negative. Two representative images showing opening or its lack are shown. **(b)** Overall, over 91.2% (31 out 34 mice) showed successful BBB opening in the targeted sites. **(c)** Examples of positive BBB opening sites for each group showing the approximate range of possible results (N=3 per group). Scale bars are 1 mm for all panels.
